# A pipeline for identifying small noncoding RNA (sRNA) candidates in bacteria

**DOI:** 10.64898/2026.07.02.735529

**Authors:** Sinda Elhedi, Kadidia Dite Selly N’Diaye, Jonathan Perreault

## Abstract

Bacterial small non-coding RNAs (sRNAs) are central post-transcriptional regulators, yet their computational identification suffers from high false-positive rates due to transcriptional noise and the absence of canonical coding features.

We developed a three-stage pipeline integrating sRNA prediction (sRNA-Detect), transcription start site mapping (TSSAR, dRNA-seq), and Rho-independent terminator detection (RNIE), applied across nine phylogenetically diverse bacterial species spanning six phyla.

Sequential filtering achieved 1.4 to 33 fold precision improvements across nine species, reducing candidate sets by up to 99.6% while recovering known sRNAs at rates reflecting reference database depth (6% recall in S. aureus, 33-34% in E. coli and S. enterica)

TSS and RIT constraints constitute universal, genome-size-independent biological filters that substantially enrich sRNA predictions across bacterial diversity. Precision variation across species reflects database incompleteness rather than pipeline failure, with unmatched predictions in poorly annotated organisms representing candidate novel sRNAs rather than false positives. RNA-seq coverage depth provides a reliable secondary indicator of biological relevance, though its interpretation requires accounting for sequencing depth variation across datasets.

## Introduction

Small RNAs (sRNAs) are non-coding RNA molecules of 20-500 nucleotides that serve as central post transcriptional regulators of bacterial physiology. Through base-pairing interactions with target mRNAs, these molecules control diverse cellular processes including stress adaptation, quorum sensing, and virulence factor expression (Storz et al., 2011, Gottesman & Storz, 2011). Their regulatory importance extends to iron homeostasis and antimicrobial resistance processes essential for bacterial survival in host environments (Vanderpool & Gottesman, 2004; Massé & Gottesman, 2002).

Bacterial sRNAs operate through two mechanistically distinct pathways. Cis-acting sRNAs are transcribed antisense to their target genes, exhibiting near-perfect complementarity that enables precise, gene-specific regulation through direct base-pairing. This architecture allows rapid response to environmental changes with minimal regulatory complexity. In contrast, trans-acting sRNAs can originate from distant genomic loci and typically display limited complementarity with their targets, often requiring the RNA chaperone Hfq to facilitate and stabilize regulatory interactions (Vogel & Luisi, 2011). Trans-acting sRNAs frequently coordinate multi-target regulons, enabling synchronized cellular responses to environmental fluctuations.

Despite extensive characterization in model organisms such as *Escherichia coli* and *Salmonella enterica* where 98 and 118 sRNAs have been experimentally validated, respectively, as of 2022 (Boutet et al. 2022), the vast majority of bacterial species remain poorly explored with regards to their sRNAs. This knowledge gap stems from limited transcriptomic coverage and the absence of specialized computational frameworks adapted for taxonomically diverse bacteria. Addressing this disparity is critical for understanding bacterial adaptation, pathogenesis, and metabolic versatility across ecological niches.

The widespread adoption of high-throughput RNA sequencing (RNA-seq) has transformed bacterial transcriptomics, enabling genome-wide sRNA discovery across numerous species (Wang et al., 2009). Computational tools such as sRNA-Detect leverage RNA-seq data to identify candidate sRNAs based on expression profiles, coverage uniformity, size constraints, and intergenic localization (Peña-Castillo et al., 2016). However, these expression-based approaches consistently suffer from elevated false-positive rates due to several inherent limitations:

Transcriptional noise: RNA-seq captures all transcribed regions, including transient transcripts, degradation intermediates, and antisense transcription resulting from promoter read-through or spurious initiation.

mRNA processing artifacts: Cleaved mRNA fragments can mimic authentic sRNAs in both size distribution and coverage patterns, particularly when derived from abundant transcripts.

Functional ambiguity: Expression-based methods cannot distinguish between functional regulatory RNAs and non-functional transcriptional byproducts without additional biological context.

Antisense contamination: Many predicted antisense sRNAs may represent transcriptional noise rather than bona fide regulatory elements with characterized functions; or novel functional antisense RNAs.

The absence of transcriptional context, specifically transcription start sites (TSSs) and termination signals, makes it difficult to distinguish genuine regulatory sRNAs from spurious transcripts. This limitation has profound consequences for experimental validation efforts, as researchers may expend considerable resources characterizing non-functional candidates.

We hypothesized that integrating multiple layers of transcriptional evidence would substantially reduce false positives while maintaining sensitivity for genuine sRNAs. This approach exploits fundamental biological principles: most authentic sRNAs exhibit discrete TSSs indicating independent transcriptional units and intrinsic terminators ensuring proper 3′ end formation (Jackson et al.2017).

Transcription start site mapping through differential RNA-seq (dRNA-seq) provides single-nucleotide resolution of transcription initiation by exploiting the differential sensitivity of primary (5′-triphosphate) versus processed (5′-monophosphate or 5’ hydroxyl) transcripts to 5′-terminator exonuclease (TEX) treatment (sharma et al. 2010).

Rho-independent terminators (RITs) represent the predominant transcription termination mechanism in bacteria. These structures consist of GC-rich hairpin loops followed by U-rich tracts that destabilize the RNA-DNA hybrid, triggering RNA polymerase pausing and transcript release (Gusarov & Nudler 1999; d’Aubenton et al. 1991). Computational detection through covariance models implemented in RNIE (Gardner et al. 2011) allows homology searches independent of primary sequence conservation, as terminator structures exhibit characteristic architectural features recognizable across substantial phylogenetic distances. The presence of canonical terminators downstream of predicted sRNAs provides strong support for authentic transcriptional units with defined 3′ boundaries (Brenes-Álvarez et al. 2016).

We present a systematic evaluation of integrative sRNA prediction across nine phylogenetically diverse bacterial species: the Gram-positive models *Staphylococcus aureus* USA300, *Bacillus amyloliquefaciens* FZB42, and the clinically critical nosocomial pathogen *Clostridium difficile* 630; as well as the epsilon-proteobacteria *Helicobacter pylori* 26695 and *Campylobacter jejuni* 81116; the gamma-proteobacterial model organisms *Escherichia coli* K-12 MG1655 and *Salmonella enterica* ST4/74; and the environmentally significant species *Shewanella oneidensis* MR-1 and *Methylorubrum extorquens* DM4. This phylogenetic breadth enables assessment of pipeline performance across diverse genomes, regulatory strategies, and database coverage levels.

By systematically filtering sRNA predictions through TSS proximity requirements and terminator presence, we demonstrate substantial false-positive reduction while retaining experimentally validated candidates. Our specific objectives were to:

Develop a robust computational framework integrating transcriptional context to reduce false-positive sRNA predictions

Benchmark the approach on well-characterized organisms to validate effectiveness

Apply the pipeline to less-characterized bacterial species to demonstrate broad applicability Quantify precision improvements achieved through systematic filtering

Provide a flexible framework adaptable for diverse bacterial taxa and experimental conditions

## Materials and Methods

Figure 1, presents an overview of the proposed bioinformatics pipeline, illustrating the main stages involved in the identification and validation of small non-coding RNAs (sRNAs).

**Figure 1:**
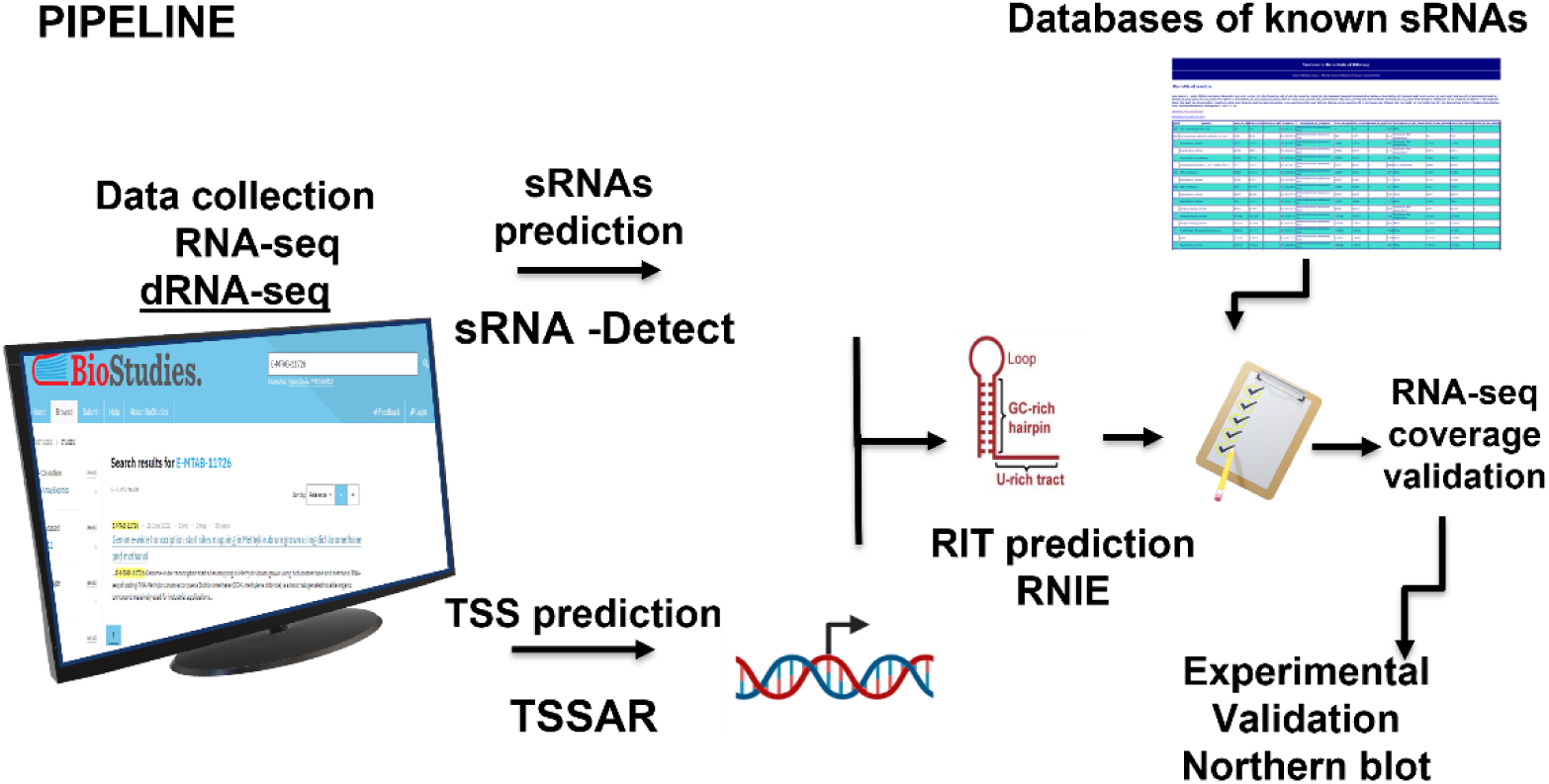
Pipeline presentation The workflow begins with the retrieval of datasets from publicly available repositories followed by sequence alignment against the target genome. Candidate sRNAs are then refined through the identification of transcriptional features, including transcription start sites (TSSs) by TSSAR and rho-independent terminators (RITs) by RNIE. Subsequent filtering steps are applied to retain high-confidence candidates, resulting in a final catalog of predicted sRNAs for downstream analyses and validation.

### Data collection and species selection

We implemented a tiered validation strategy utilizing both well-characterized benchmark species and less-studied bacteria. This approach allowed validation against known sRNA repertoires while demonstrating applicability to understudied organisms.

Nine phylogenetically diverse bacterial species were selected, spanning six phyla: three benchmark species with well-characterized sRNA repertoires (*Escherichia coli* K-12 MG1655; *Staphylococcus aureus* USA300; *Salmonella enterica* Typhimurium ST4/74) and six test species representing understudied diversity *(Helicobacter pylori* 26695; *Campylobacter jejuni* 81116*; Methylorubrum extorquens* DM4; *Shewanella oneidensis* MR-1; *Clostridium difficile* 630Δerm; *Bacillus amyloliquefaciens* FZB42). Publicly available RNA-seq and differential RNA-seq datasets were sourced from BioProjects (Table S1).

### Read alignment

RNA-seq reads for each species were aligned to their respective reference genomes using HISAT2 version 2.2.1 (Kim et al. 2015) with default parameters. Alignment files were converted to BAM format and sorted using SAMtools version 1.15 (Li et al. 2009).

### Transcription Start Site Identification Using TSSAR

Differential RNA-seq (dRNA-seq) exploits the differential sensitivity of RNA 5′ ends to 5′-terminator exonuclease (TEX) treatment. Primary transcripts retain 5′-triphosphate ends and resist degradation, whereas processed transcripts with 5′-monophosphate (and 5’-OH) ends are selectively degraded. By comparing TEX-treated (+TEX) and untreated (−TEX) libraries, dRNA-seq enables precise genome-wide identification of transcription start sites (Sharma et al. 2010).

### TSSAR Implementation

TSSAR (Transcription Start Site Annotation Regime) (Sharma et al. 2010) identifies TSSs by modeling read start count differences between +TEX and −TEX libraries using Poisson and Skellam distributions. TSSAR assigns a differential score to each genomic position, reflecting the statistical significance of enrichment in the +TEX library.

#### Key parameters

The command line used to perform TSSAR analysis of dRNA-seq data was:

. /TSSAR --libP I<libraryP.sam> --libM I<libraryM.sam> [--score I<p|d>] [--fasta I<genome.fa>] [--minPeak I<INT>] [--winSize I<INT>]

Where:

--minPeak 3: Minimum read count required for TSS calling

--score d: Differential scoring mode between +TEX and −TEX libraries

--winSize 1000: Window size for peak scanning and background estimation

We evaluated 13 discrete thresholds spanning four orders of magnitude (0 to 40,000) to capture the full dynamic range of TSS statistical confidence. This range encompasses low-confidence predictions (score < 100), moderate confidence suitable for exploratory analysis (100-1000), high confidence (1000-2000), and extremely stringent thresholds for mechanistic studies (>2000). The geometric progression enables systematic evaluation of precision-sensitivity trade-offs across the confidence spectrum.

### sRNAs candidate prediction using sRNA-Detect

#### Algorithm Principles

sRNA-Detect (Peña-Castillo et al., 2016), identifies small RNA candidates based on uniform coverage patterns derived from RNA-seq read alignment data. The algorithm operates on the principle that authentic small transcripts should exhibit consistent read coverage across their entire length, distinguishing them from fragmented mRNA degradation products or spurious transcription.

##### Key Parameters

The command line used to perform sRNA-Detect analysis of RNA-seq data was:

nextflow detect_filter_sRNA.nf --alignmentDir --annotatedGenomeFile --outputDir –output --idPrefix RCAP_rcs --minLength 20 --maxLength 500 --minHeight 10

--minLength 20: Minimum transcript length (nt)

--maxLength 500: Maximum transcript length (nt)

We also evaluated 13 discrete score thresholds to assess their impact on prediction precision. Higher thresholds retain only candidates with highly uniform coverage and strong expression, while lower thresholds capture candidates with moderate expression levels.

The initial sRNA candidates considered for the rest of the analysis are not raw tool outputs, but a pre-filtered set (sRNA-Detect separate and filter annotated RNA): predictions that overlap previously annotated features (rRNA,tRNA,tmRNA,RNase_P_RNA,protein_coding) in the corresponding GFF file were not retained.

### Integration strategy: TSS-sRNA Intersection analysis

Predicted sRNAs candidates were intersected with TSS predictions using bedtools closest (Quinlan & Hall, 2010) with strand-specificity enabled and distance reporting.

#### Key parameters

The command line used to perform TSS intersection to predicted sRNAs was:

bedtools closest -a <FILE> -b <FILE> -s -D ref

-s: Strand-specificity

-D ref: Report distance with respect to the reference genome

A critical pipeline parameter is the TSS distance threshold, defining the maximum allowable distance between an sRNAs 5′ start site and a TSS position. For this, we used a “TSS distance threshold” of −60 bp upstream and +20 bp, downstream applied uniformly across all species. The threshold (−60 bp upstream, +20 bp downstream) is biologically motivated: Reverse Transcription (RT) performed on RNA prior to sequencing may not always reach the 5′ end of the RNA, hence a margin of −60 bp to account for this; and in principle we do not expect any RT to go beyond the TSS, but to account for technical variability in TSS mapping precision due to library complexity or read depth and potential annotation uncertainties, a margin of +20 bp is accepted. Only sRNAs with at least one TSS with a position that fell within −60 bp upstream to +20 bp downstream of the predicted 5′ start position of the candidate sRNA (on the same genomic strand) were retained for downstream analysis.

### RNIE Implementation

RNIE (Gardner et al. 2011) uses covariance models (CMs) to predict Rho-independent terminators short genomic sequences characterized by G+C-rich hairpin loops followed by a polyuridine tail that cause bacterial RNA polymerase to pause and terminate transcription.

#### Two Operating Modes

##### Genome Mode

Optimized for high-throughput genome annotation with rapid scanning (∼43 kb/s), very low false positive rate (∼1.7 FP/M), sensitivity of 0.70,

##### Gene Mode

Optimized for individually annotating downstream regions of genes with higher sensitivity (0.83) but slower scanning (∼1 kb/s) and higher false positive rate (∼9.6 FP/MB)

#### Implementation parameters

TSS-associated sRNA intervals were extended 150 nucleotides downstream to capture potential terminator regions.RNIE was executed using built-in covariance model: erpin-rho.cm

#### Key parameters

The command line used to perform an RNIE analysis on intersected TSS candidates was:

perl rnie.pl -m erpin-rho.cm -f <sequences< -g –sensitive (sensitive for gene mode)

Parameters:

-m: Covariance model

-g: Produce output in GFF format

--sensitive|--gene: Run in targeted-region mode. Uses a parameter set tuned for high sensitivity. Primarily for answering the question: "Could there be a Rho independent terminator following my gene of interest?"

Each candidate region received a bit score quantifying similarity to canonical RIT models. Sequences were extracted using bedtools getfasta with strand-specificity maintained. For RIT-associated sRNAs, final coordinates combined the original sRNA 5′ start with the detected terminator 3′ end, creating complete transcriptional units with defined boundaries.

Initial predictions employed cmsearch (embedded in RNIE) with default bit score threshold (-T parameter) optimized for reference organisms. However, since our dataset comprises phylogenetically diverse bacterial species that may not have been fully represented in the benchmark sets used to calibrate cmsearch defaults, we assessed whether adjusting the bit score threshold could improve prediction recall without introducing excessive false positives. Notably, preliminary validation studies in our laboratory identified a novel sRNA (sRNA 1969; by Emillie Boutet, 2023) that was not detected using the default threshold parameters. We therefore re-ran predictions with the bit score threshold (-T parameter) adjusted to 4 across all species, a more permissive threshold calibrated to enhance sensitivity for non-benchmark organisms and capture previously unidentified sRNAs such as sRNA 1969.

### Database Sources

sRNA sequences were validated to database sources by species. Moreover, sRNAs from Rfam were also used.

### Test species

#### *E. coli* K-12 MG1655

A list containing a total of 116 *E. coli* sRNAs combined from BSRD and Rfam (Li et al. 2013, Kalvari et al., 2021)

#### *Salmonella enterica* ST4/74

A list containing 143 *S.enterica* sRNAs from BSRD and Rfam (Li et al. 2013, Kalvari et al., 2021)

#### *Staphylococcus aureus* USA300

SRD (Staphylococcal Regulatory RNAs Database) (Sassi et al. 2015) with 88 distinct sRNA entries covering strain USA300_FPR3757

### Less characterized species

#### *Bacillus amyloliquefaciens* FZB42

BSRD containing 28 *Bacillus* spp sRNAs

#### *Helicobacter pylori* 26695

BSRD containing 80 *Helicobacter* spp. sRNAs +1 sRNA from Rfam (Li et al. 2013, Kalvari et al., 2021).

#### *Campylobacter jejuni* 81116

Manually curated database from Dugar et al,.2013 study containing 21 experimentally characterized *C. jejuni* sRNAs by Northern blot (Table S3)

#### *Shewanella oneidensis* MR-1

Rfam and NCBI annotations sRNAs (17 sRNAs)

#### *Methylorubrum extorquens* DM4

RiboGap database (Naghdi et al. 2017) containing 7 characterized *Methylorubrum* sRNAs.

#### *Clostridium difficile* 630

Rfam and experimentally validated (Northern blot) predicted sRNAs from a study by Soutourina et al. 2020 results in 14 sRNAs.

### BLAST Implementation

Sequences were blasted to known sRNAs databases through the command line

blastn -task blastn-short -query <sequences> -db <database> -outfmt 6 -evalue 1e-1 - perc_identity 95

#### Key parameters

blastn-short is optimized for the alignment of short nucleotide sequences, such as small RNAs, by using a reduced word size and adjusted scoring parameters. It increases sensitivity for detecting matches of short sequences while maintaining specificity

To reduce weak and potentially spurious matches, a bit score filter (> 60) was applied to all BLAST results, ensuring that only alignments with significant sequence homology were considered. (nonfiltered results are presented in Table S4)

#### Threshold Optimization and Statistical Analysis

We implemented a comprehensive grid search approach, evaluating all combinations of predicted sRNAs and TSS scores across 13 thresholds, resulting in 13 × 13 = 169 combinations per species. Additionally, we analyzed sRNA predictions without TSS filtering to establish baseline performance.

#### For each parameter combination, we computed

- Number of TSS predictions retained at given TSS score
- Number of sRNA predictions retained at given sRNA score
- Number of sRNAs-TSS intersections
- Number of BLAST-validated candidates (intersections with database homology for known sRNAs)
- Number of sRNAs with RIT structures detected downstream
- Number of BLAST-validated sRNAs with RIT structure

Detailed results are provided in Table S2.

Precision and Recall Calculation:

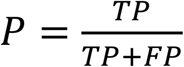, where:
P (precision) = proportion of predicted sRNAs that are correct
TP (True Positives) = predicted sRNAs that match known sRNAs
FP (False Positives) = predicted sRNAs that do not match known sRNAs
TP + FP = total number of predicted sRNAs

If precision is known:

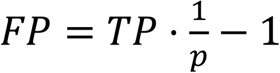 where FP=False Positives

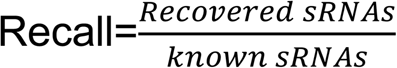

where:

Recovered sRNAs = known sRNAs correctly predicted

Known sRNAs = total number of annotated/expected sRNAs

## Results

### Pipeline overview and Recovery Strategy

Our pipeline proceeds through three sequential filtering steps (Figure 2). First sRNA candidates are predicted from RNA-seq data using sRNA-Detect. Second, these predictions are filtered by proximity to experimentally mapped TSSs from TSSAR retaining only transcripts with defined transcriptional start points. Third, retained candidates are extended downstream and analyzed with RNIE to identify Rho-independent terminators. Candidates possessing both features, termed final sRNA candidates, represent our final high confidence predictions. For most datasets, the starting number of candidates is unacceptably high to explore them one by one (up to ∼30,000), but these numbers are reduced by one to two orders of magnitude by applying filters.

**Figure 2:**
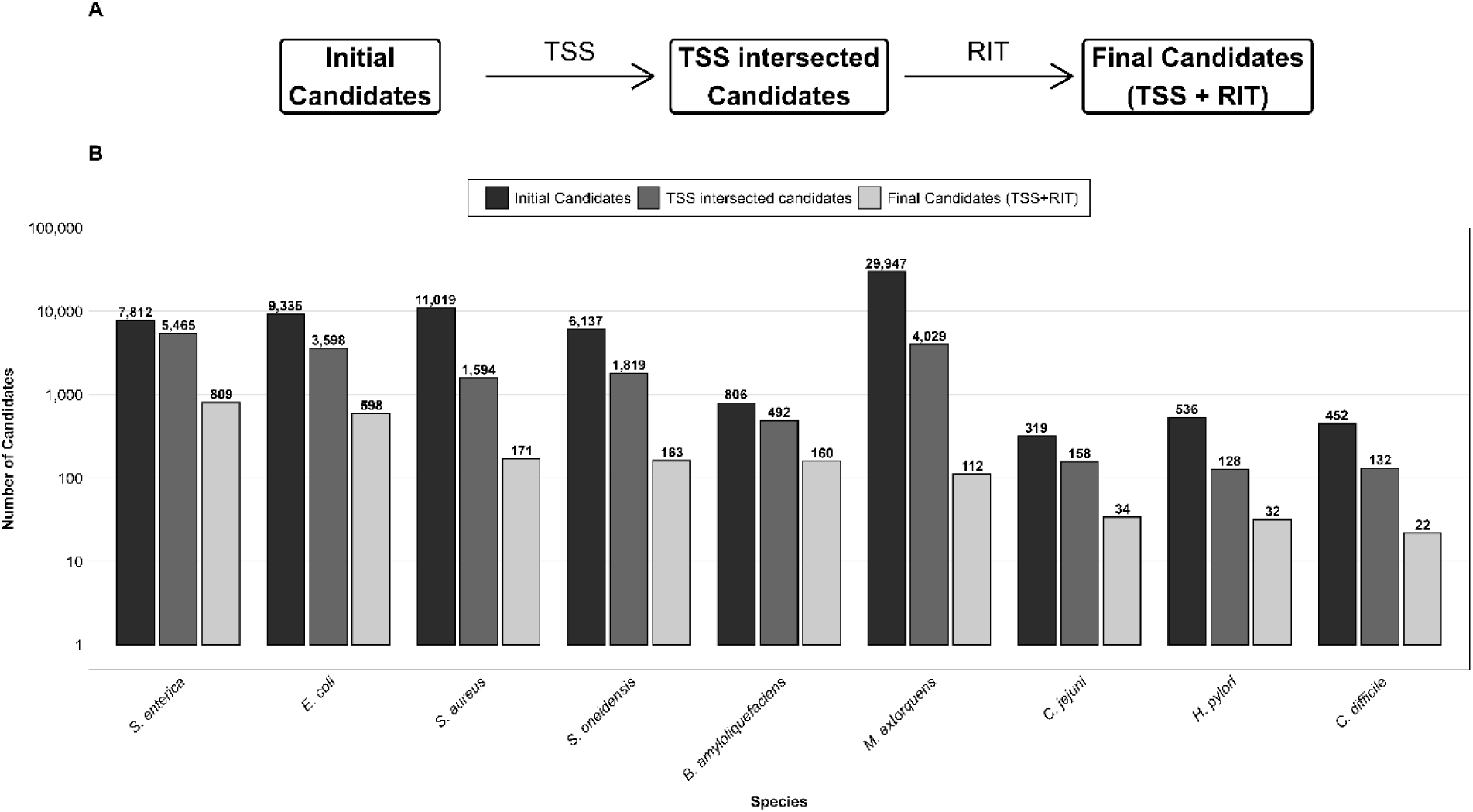
Summary of Pipeline Performance Flow diagram illustrating the three sequential filtering stages of the sRNA discovery pipeline: (B) Histogram showing the three steps progression: Initial candidates: predicted sRNAs candidates by sRNA-Detect,TSS intersected candidates : predicted sRNAs candidates intersected with TSS, Final candidates: predicted sRNAs candidates intersected with TSS and have RIT downstream.

### Overview of Pipeline Performance Across Species

To contextualize pipeline output, we distinguish two categories of species (Table S1 ).

#### Benchmark species

(*E. coli*, *S. aureus*, *S. enterica*) harbor well-documented sRNA repertoires against which precision and recall provide meaningful performance indicators. Initial candidate pools were large (7,812–11,019) with moderate-to-low baseline precision (1.2–1.9%). Sequential TSS and RIT filtering substantially enriched this output: *S. aureus* rose from 1.19% baseline to 3% precision across 171 candidates, enabled by the comprehensive SRD reference. *E. coli* reached 7% final precision while recovering 38 of 116 known sRNAs (33% recall). *S. enterica* recovered 48 of 143 reference entries (34% recall) as showed on Figure 3.

#### Test species

(*M. extorquens*, *S. oneidensis*, *B. amyloliquefaciens*, *H. pylori*, *C. jejuni*, *C. difficile*) are organisms for which the sRNA repertoire is largely uncharacterized. By this criterion, the pipeline succeeds consistently. *M. extorquens*, whose initial pool of 29,947 candidates would be impossible to validate one by one, is reduced to 112 final candidates, a 99.6% reduction while recovering 1 of 7 deposited reference sRNAs (14% recall). *S. oneidensis* is reduced from 6,137 to 163 candidates (97.3% reduction). *C. difficile* candidates are reduced from 452 to 22 candidates with 86% precision*. H. pylori* et *C. jejuni* move from 536 and 319 initial candidates respectively to 32 and 34 (22% and 9% precision respectively and reaching 14% recall for both) (Figure 3). For all test species, the filtered candidate sets constitute a prioritized, biologically motivated shortlist for experimental validation.

### Precision-Recall Trade-off

Sequential TSS and RIT filtering reduced candidate sets by 89–99% while maintaining good recall of known sRNAs (Figure 4). Among the analyzed species, *S. enterica* and *E. coli* exhibited the highest recall values, recovering approximately 34% and 33% of known sRNAs, respectively. However, their precision remained relatively low (approximately 7–8%), reflecting the large number of remaining candidates. In contrast, *C. difficile* achieved the highest precision (∼59%) while maintaining a moderate recall (∼14%), indicating highly selective candidate retention. *C. jejuni* showed intermediate performance, with a recall of approximately 14% and a precision of 9%. *H. pylori* displayed the lowest recall (∼7%) but achieved a relatively high precision (∼22%), suggesting a conservative filtering strategy that retained few but enriched candidates. *S. oneidensis* and *M. extorquens* exhibited low precision despite recalls of approximately 12% and 14%, respectively.

**Figure 3:**
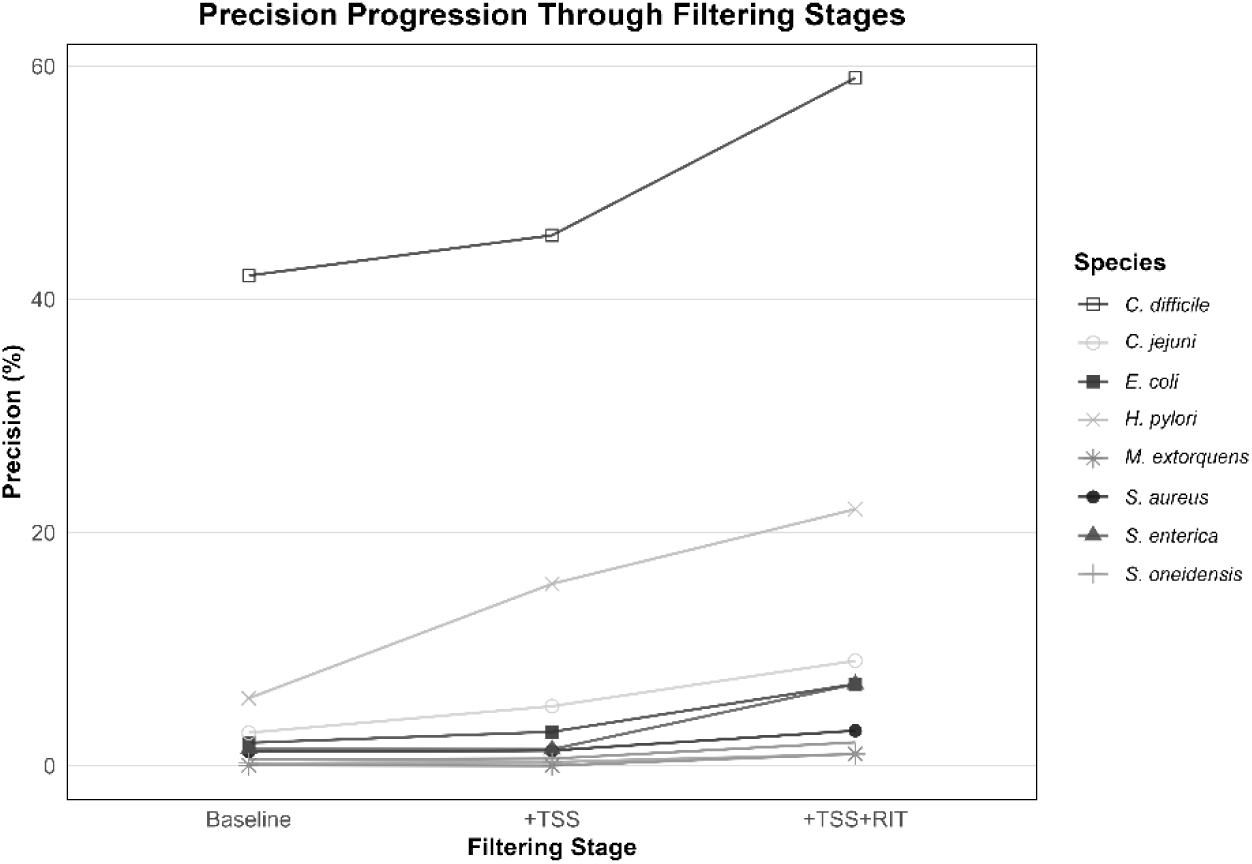
Precision Progression Across Three Filtering Stages and Nine species Precision (fraction of predictions matching known sRNAs) is shown at three successive pipeline stages: baseline sRNA-Detect predictions (Baseline), TSS intersection with baseline (+TSS), and additional RIT filtering (+TSS+RIT).

**Figure 4:**
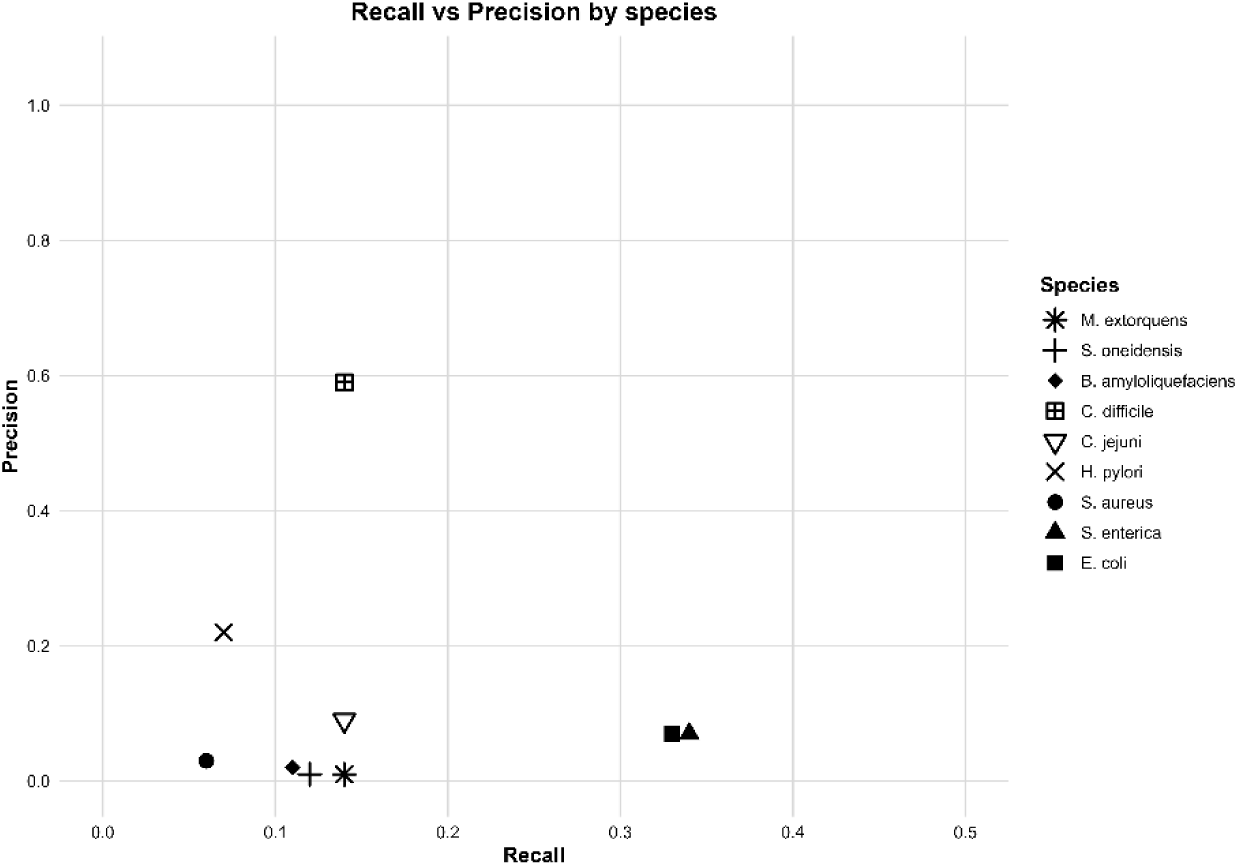
Performance Trade-offs: Recall vs Precision Recall (x-axis) = % of known sRNAs recovered. Precision (y-axis) = % of predictions homologous (at least 95%_identity) to known sRNAs in the corresponding species, as determined by BLAST.

Overall, the sequential application of TSS and RIT filters effectively reduced the candidate search space while preserving a substantial fraction of known sRNAs. The increase in precision observed in several species suggests that combining transcription start site evidence with intrinsic terminator prediction preferentially removes false-positive candidates while retaining biologically relevant sRNAs.

### Validation of final candidates with RNA-seq data

To evaluate the transcriptional support for computational predictions, final potential candidate sRNAs were assessed against RNA-seq data through read coverage analysis, whereby mean read depth was calculated for each candidate and used as a quantitative indicator of expression level. Candidates were subsequently classified into four tiers based on mean depth thresholds: highly expressed,moderately expressed,weakly expressed, and poorly expressed, reflecting candidates with insufficient RNA-seq evidence and likely representing computational artifacts.

As illustrated in Figure 5 the classification profile of validated sRNA candidates varied markedly across species, reflecting the combined influence of sequencing depth and intrinsic sRNA biology. Species datasets with the highest mean depth — *H. pylori* (2524.4 reads/position), *C.difficile (*1999 reads/position), *S. enterica* (287.7 reads/position), and *E. coli* (354.9 reads/position) — displayed bars dominated by highly expressed sRNAs classification, demonstrating that deep coverage enables consistent read accumulation above the ≥50-read threshold required for high-confidence assignment. In contrast, medium-depth dataset as *C.Jejuni* (156.7 reads/position) and low-depth datasets such as for *Shewanella* (32.3 reads/position) and *Methylorubrum* (41 reads/position) exhibited proportionally large weak fractions, suggesting that genuine sRNA transcripts were present but insufficiently covered to surpass classification thresholds, reflecting a depth artifact rather than true absence of expression.

**Figure 5:**
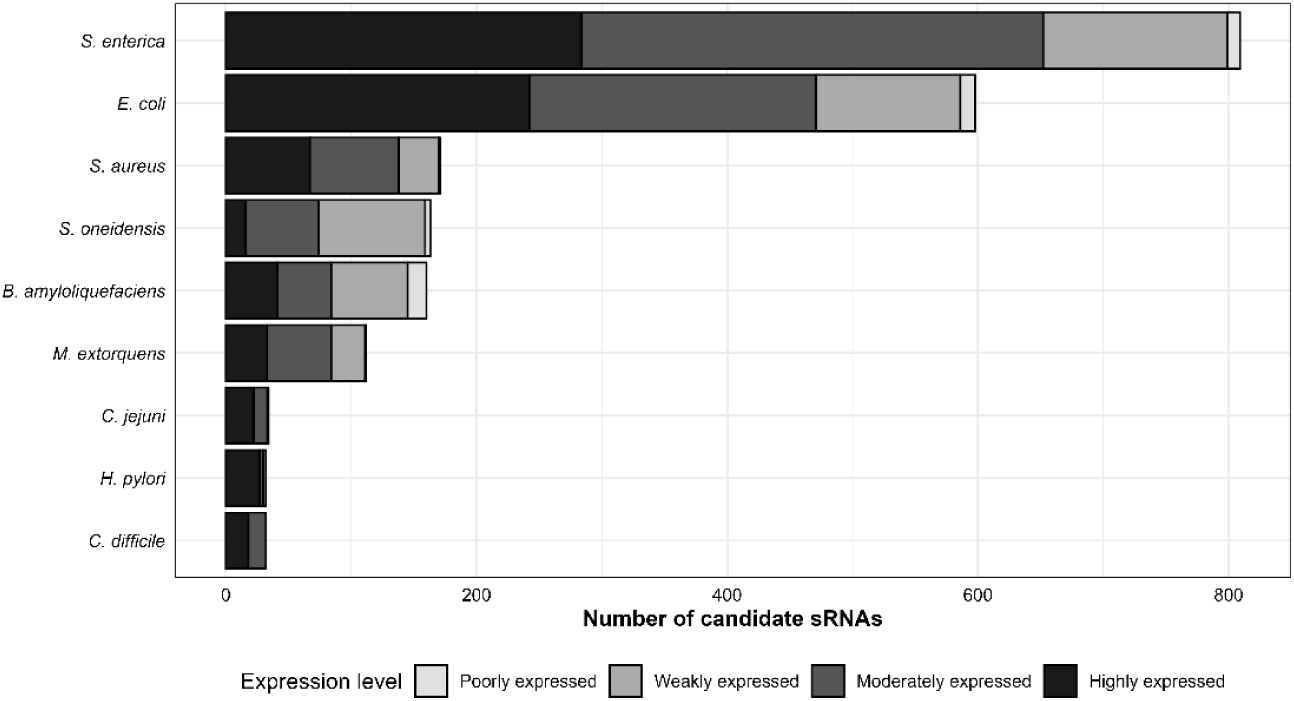
sRNAs classification by RNA-seq coverage depth across nine bacterial species Distribution of sRNA candidates across four confidence tiers based on mean read depth: Highly expressed (≥50 reads/position, black), moderately expressed (10–50 reads, medium-dark grey), weakly expressed (2–10 reads, medium-light grey), and poorly expressed (<2 reads, near-white, grey). Each bar represents the total number of candidates per species (after applying TSS and RIT filters); colors indicate the proportion in each confidence tier. Species are ordered by decreasing total candidate count from top to bottom.

To evaluate the trade-off between prediction accuracy and candidate retention, computational predictions were subjected to progressively stringent filtering at score thresholds ranging from 0 to 40,000 (Figure S1).

To be mentioned, restricting the analysis to high-scoring predictions from the outset would have prematurely eliminated candidates before transcriptional unit filtering could be applied. Valid sRNAs are supported by RNA-seq coverage at the validation stage.

In *Figure 6.A*, *E.coli* demonstrates a progressive, reduction in candidate numbers with increasing score threshold, declining from 598 candidates at threshold 0 to 34 candidates at threshold 2000 (a 94% reduction). Crucially, precision improves steadily across the entire threshold range, rising from 7 % at score 0 to 29% at score 2000. This consistent improvement indicates that higher-scoring predictions are systematically more likely to represent genuinely annotated sRNAs, suggesting the scoring algorithm effectively captures features that correlate with true sRNA biology. Conversely, recall is reduced gradually from one stringency level to the next, declining gradually from 33% at threshold 0, to ∼ 9 % at threshold 2000.

**Figure 6:**
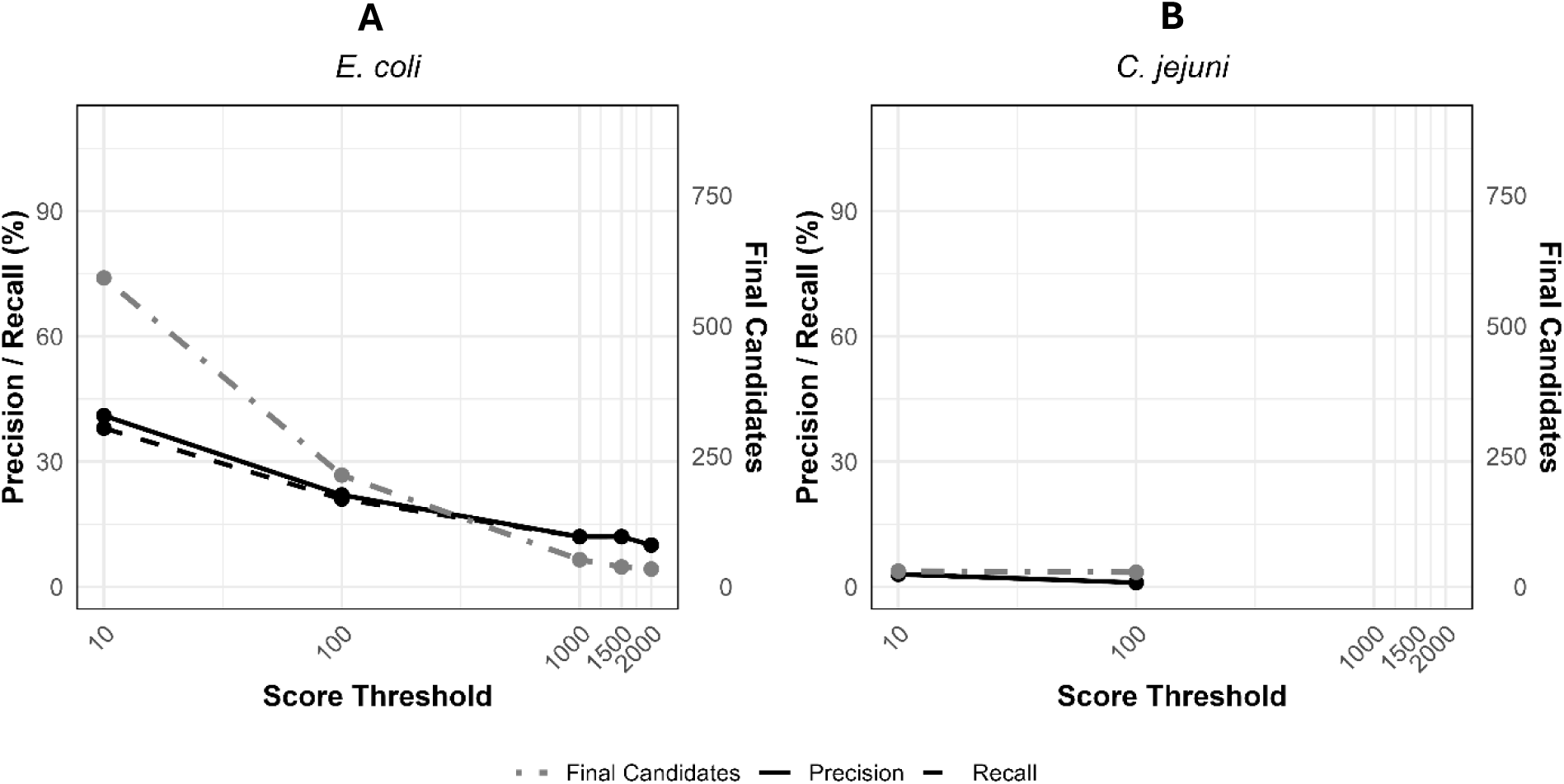
Effect of score threshold on final sRNAs Discovery: well-Annotated vs poorly annotated species Comparison of *E. coli* K-12 (A) and *C. jejuni* (B), for final numbers of retained candidates, precision and % recall of known sRNAs within these respective species.(plots of all species are presented in Figure S2 in Supplementary materials).

In stark contrast, *C.jeju*ni (Figure 6.B) presents a markedly different filtering response, exemplifying prediction challenges in organisms with limited functional annotation and lower RNA-seq coverage depth. The candidate pool undergoes depletion at higher thresholds: declining from 34 candidates at score 0 to zero candidates by score 1500. However, recall drops precipitously from 14% at threshold 0 to 4.5 % at score 100 and reaches zero by score 1500.

This collapse in recall demonstrates that aggressive filtering eliminates many true sRNAs from the candidate set, despite improving precision metrics.

These observations and all species comparison (Figure 7) justify adopting a score threshold of 0 in the main pipeline analysis across all species, reducing the risk that potentially meaningful candidates are prematurely discarded.

**Figure 7:**
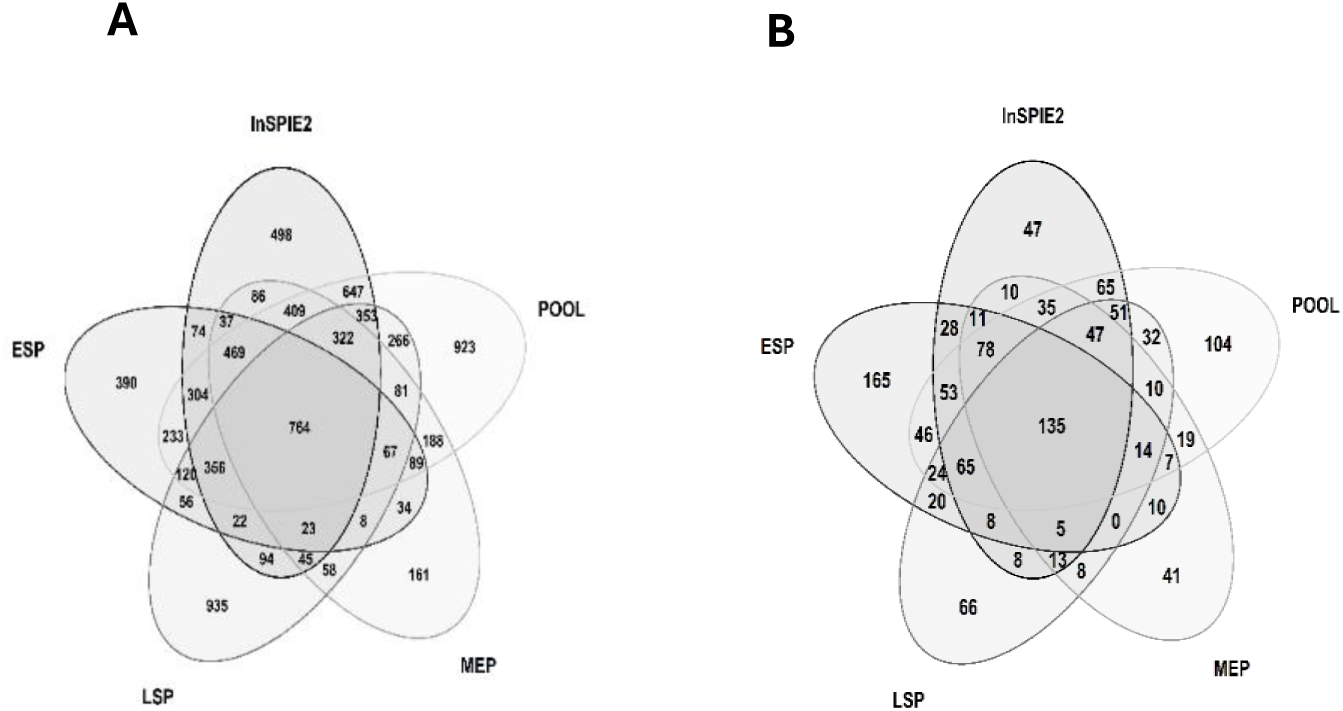
Condition-Dependent sRNA Expression in S. enterica ST4/74 Two complementary Venn diagrams comparing sRNA detection across five experimental datasets in Salmonella enterica serovar Typhimurium ST4/74. (A) Raw sRNA-Detect predictions before filtering. (B) Pipeline-filtered candidates (TSS + RIT). Five datasets profiled: MEP (mid-exponential phase), ESP (early stationary phase), LSP (late stationary phase), InSPI2 (SPI-2-inducing conditions), and POOL (22 combined environmental conditions (Kröger et al,2013).

**Figure 8:**
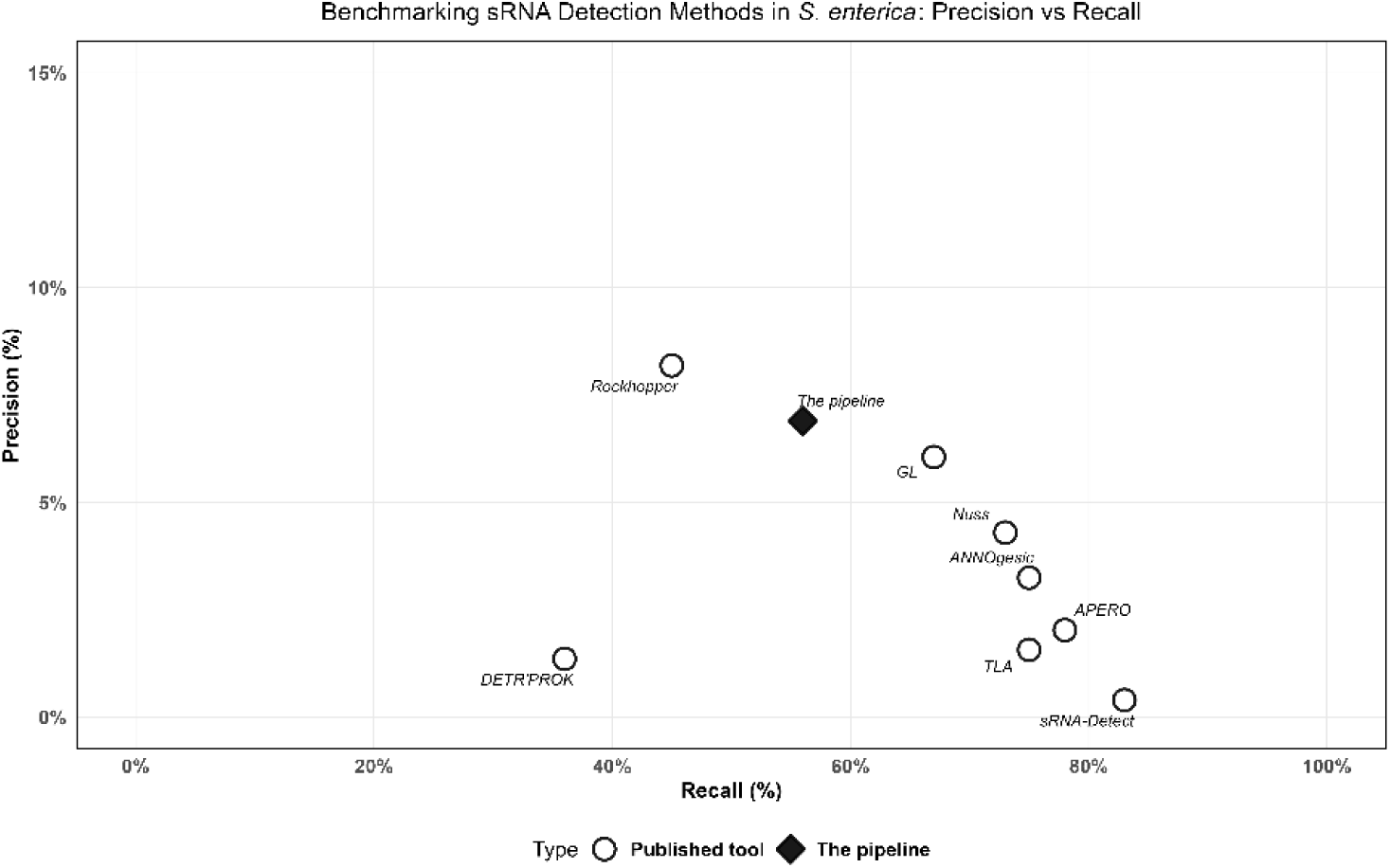
Benchmarking sRNA Detection Methods in S. enterica — Precision versus Recall Precision and recall performance of eight published sRNA detection pipelines against this study’s approach. Each point represents one method’s operating point. Symbols: (white circle) = published tools; (black diamond) = this study. Published tools include ANNOgesic, APERO, DETR’PROK, GL, Nuss, TLA, Rockhopper, and sRNA-Detect. This study shows post filtering results.Precision for this plot is calculated by P=Models/Total predicted similarly to the study.

Condition-dependent transcription shapes sRNA discovery in *Salmonella enterica* case To evaluate differential sRNA expression across growth conditions, RNA-seq data from

*Salmonella enterica* serovar Typhimurium ST4/74 were analyzed across five experimental samples from BioProject PRJNA215033 (Kröger et al., 2013): mid-exponential phase (MEP), early stationary phase (ESP), late stationary phase (LSP), SPI-2-inducing conditions (InSPI2), and a pooled sample derived from 22 diverse environmental conditions (POOL).The pipeline was executed independently for each condition to capture condition-specific expression patterns.

Across all conditions, 1225 unique loci were identified, underscoring the complexity and plasticity of the sRNA landscape. Among these, only 135 candidates were consistently detected in all conditions, defining a relatively small core set of constitutively expressed sRNAs. In contrast, 423 loci were strictly condition-specific, while the remaining candidates were shared across subsets of conditions, indicating graded or context-dependent expression patterns. The number of detected loci varied markedly between conditions, with POOL yielding the highest count (785), followed by ESP (669), InSPI2 (659), LSP (506), and MEP (443) as shown in Figure 7. However, the high yield observed in POOL primarily reflects the aggregation of signals from multiple environments rather than an increase in biologically coherent transcriptional programs.

A more informative pattern emerges when focusing on condition-specific sRNAs post filtering. ESP exhibited the highest number of unique candidates (165), substantially exceeding other conditions, including POOL (104), LSP (66), InSPI2 (47), and MEP (41) (Figure 7)

Importantly, the pooled dataset, while increasing overall detection sensitivity, appears to dilute condition specificity by merging distinct transcriptional states. As a result, sRNAs detected in POOL may represent cumulative expression rather than condition-resolved regulation.

These observations closely align with the previously reported distribution of condition-specific transcription start sites (TSS) in the same study (Kröger et al., 2013), reinforcing the idea that transcription initiation is a primary determinant of sRNA emergence. Conditions enriched in unique TSS, such as ESP (34 candidates), tend to produce a higher number of specific sRNAs.

Together, these results highlight a fundamental trade-off in sRNA discovery: while pooled datasets maximize sensitivity and broaden the detectable repertoire, analyses of individual conditions are essential to resolve regulatory specificity and uncover biologically meaningful sRNA expression patterns.

### Positioning of the pipeline relative to existing tools (*S. enterica* case study)

For *Salmonella enterica*, several sRNA prediction tools have been evaluated in the literature by Leonard et al. 2019, including ANNOgesic, APERO, DETR’PROK, GL, Nuss, TLA, Rockhopper, and sRNA-Detect.

These approaches exhibit different trade-offs between recall and specificity: the best tools achieve a recall of approximately 0.75–0.83 on their reference sRNA sets, but at the cost of hundreds to several thousands of predicted transcripts per species. For example, sRNA-Detect alone predicts 17,662 transcripts in *S. enterica*, of which 71 correspond to known sRNAs, resulting in a substantial burden of manual screening and experimental validation.

In this context, our multi-layered pipeline, applied to *S. enterica*, recovers 47 known sRNAs out of 84 models reported in Leonard et al. 2019, while reducing the final candidate set to 809 regions. In other words, it maintains a recall of 0.5 on known sRNAs while decreasing by nearly 100× the number of transcripts that need to be examined compared with raw sRNA-Detect output. This combination of good recall with a restricted number of candidates shows that the systematic integration of TSS and Rho-independent terminators can turn a sensitive tool with limited specificity such as sRNA-Detect into a much more specific and experimentally tractable pipeline.

## Discussion

### From raw predictions to significant candidates

Expression-based tools like sRNA-Detect excel at sensitivity but struggle with specificity, predicting thousands of short transcripts where false positives scale dramatically with genome size. In the genome of *S.oneidensis* (4.97 Mb), 99% of predictions are noise; for *S.enterica* (4.88 Mb), false-positive rates reach 88%. Rather than tuning statistical thresholds, our pipeline imposes two universal biological rules grounded in transcriptional mechanics: candidates must overlap experimentally mapped transcription start sites (TSS), because independent genes begin at promoters; and possess Rho-independent terminators (RIT), because functional transcripts require defined 3′ boundaries.

This sequential filtering delivered robust precision gains across nine phylogenetically diverse species, but the magnitude and interpretation of these gains differ fundamentally between two groups of organisms: those with large initial prediction sets (*S. aureus*: 11,019; *E. coli*: 9,335; *S. enterica*: 7,812; *M. extorquen*s: 29,947; *S. oneidensis*: 6,137) where baseline precision is extremely low (0.03–1.96%), and those where sRNA-Detect itself generates compact prediction sets (*H. pylor*i: 536; *C. jejuni:* 319; *C. difficile*: 452) yielding comparatively higher baseline precision (2.82–42.04%). This distinction is not incidental — it reflects fundamental differences in genome-wide transcriptional complexity and the density of spurious transcriptional signals detectable by expression-based tools. In both groups, however, TSS and RIT filtering consistently enriched true positives and reduced candidate burden.

### Large prediction set species — dramatic noise reduction

For organisms where sRNA-Detect generates thousands to tens of thousands of candidates, baseline precision is necessarily low because the prediction space vastly exceeds the known sRNA complement. *Staphylococcus aureus* USA300 exemplifies this: 11,019 initial candidates yield only 1.19% baseline precision, yet sequential filtering reduces this to 171 final candidates (98.4% reduction) while recovering 5 of 88 known sRNAs (6% recall) at 3% final precision — a 2.5-fold improvement. Similarly, *M. extorquens* DM4 begins with 29,947 predictions (0.03% baseline precision), the largest initial set across all nine species, reflecting the combination of a large genome (7.0 Mb) and limited transcriptomic annotation; filtering reduces this to 112 candidates (99.6% reduction), recovering 1 of 7 known sRNAs (14% recall) at 1% final precision — a 33-fold improvement. *E. coli* K-12 MG1655 and *Salmonella enterica* ST4/74 follow the same pattern: from 9,335 and 7,812 initial predictions at 1.96% and 1.42% baseline precision respectively, filtering yields 598 and 809 final candidates at 7% final precision, recovering 38 of 116 (33% recall) and 48 of 143 (34% recall) known sRNAs. *Shewanella oneidensis* MR-1 and *Bacillus amyloliquefaciens* FZB42 similarly benefit, with candidate reductions exceeding 97% and final precisions of 1–2%.

### Compact prediction set species — precision already elevated at baseline

In contrast, *H. pylori*, *C. jejuni*, and *C. difficile* produce far fewer initial candidates (536, 319, and 452 respectively), which directly explains their higher baseline precision relative to the large-set species. This is not evidence of superior sRNA-Detect performance in these organisms per se, but rather reflects reduced transcriptional complexity, smaller genome sizes (1.67, 1.64, and 4.11 Mb respectively), or a lower density of spurious transcriptional signals detectable under the profiled conditions. *Clostridioides difficile* achieves the highest baseline precision of any species at 42.04% (190 of 452 initial predictions matching known sRNAs), rising to 59% after sequential filtering across 22 final candidates — a 1.4-fold improvement recovering 2 of 14 known sRNAs (14% recall). *Helicobacter pylori* reaches 22% final precision across 32 RIT-filtered candidates, a 3.8-fold improvement from 5.78% baseline, recovering 6 of 81 known sRNAs (7% recall).

*Campylobacter jejuni* achieves 9% final precision across 34 candidates, a 3.2-fold improvement from 2.82% baseline, recovering 3 of 22 known sRNAs (14% recall). Importantly, while these species show smaller absolute fold-improvements, this reflects their already-elevated baselines rather than reduced pipeline effectiveness.

### Database reference and Performance Gaps

Recall varied substantially across all nine species, and this variation reflects annotation depth far more than pipeline performance. The database landscape varies enormously: *E. coli* K-12 has 116 validated sRNAs; *S. enterica*, 143 entries; *S. aureus*, 88 distinct entries from the SRD (Staphylococcal Regulatory RNAs Database); and *H. pylori,* 81 BSRD entries. These organisms have been subjected to decades of intensive transcriptomic investigation, including multiple rounds of RNA-seq, dRNA-seq, and experimental validation (Storz et al. 2011, Sharma et al. 2010), providing reference sets comprehensive enough to meaningfully benchmark recall.

In stark contrast, *M. extorquens* DM4 has only seven characterized sRNAs deposited in RiboGap. This is not surprising: *M. extorquens* has received comparatively little transcriptomic attention, and sRNA annotation in non-model alphaproteobacteria lags far behind that of enterobacteria (Boutet et al. 2022). Genome-wide surveys of well-characterized bacteria suggest that dozens to over one hundred sRNAs are typical for organisms of comparable genome size (Sharma et al. 2010), implying that the seven deposited entries for *M. extorquens* DM4 represent a small fraction of its true sRNA repertoire. This annotation gap directly explains apparent low recall: pipeline predictions matching genuine but unannotated sRNAs inevitably register as false positives against an incomplete reference set.

Similarly, *C. jejuni* (22 sRNAs from manually curated literature), *B. amyloliquefaciens* (28 BSRD entries), and *S. oneidensis* (17 annotated sRNAs) represent understudied genomes where sRNA discovery lags. For these species, unmatched predictions represent genuine discoveries awaiting experimental validation rather than pipeline errors — an opportunity for new sRNA gene discovery that is structurally invisible to precision-based metrics.

### Capturing Divergent Terminators Through Threshold Calibration

Default RNIE (infernal covariance) models, calibrated on well-represented bacterial lineages, miss species-specific Rho-independent terminator variants. Relaxing the cmsearch score threshold from default 16 to 4 expanded the number of candidates sRNAs across all nine organisms. Critically, this relaxation recovered sRNA1969 in *M. extorquens* DM4 a sRNA previously validated experimentally in our laboratory (Boutet, 2023) but undetectable under default stringency. This recovery provides direct empirical support for the hypothesis that threshold relaxation rescues biologically real sRNAs that are systematically excluded when the detection model is applied to organism underrepresented in its training data, and that the expanded candidate pool is not composed exclusively of false positives.

The magnitude of candidate expansion varied across organisms in a manner consistent with both phylogenetic distance from the RNIE training set and nucleotide composition bias.

Expansion was most pronounced in AT-biased *H. pylori* (32-fold increase), substantial in enterobacteria like *E. coli and S. enterica* (∼ 3-fold expansion), and minimal in in C*. jejuni*, despite a comparable genome size to *H. pylori*, this may reflect a greater reliance on Rho-dependent termination in *C. jejuni*, resulting in a reduced density of Rho-independent terminator structures detectable by RNIE. While Rho-dependent termination is documented across bacteria (Ciampi 2006), genome-wide quantification of termination pathway usage in *C. jejuni* remains limited.

### Expression Depth and multi condition analysis

RNA-seq coverage depth emerged as a reliable indicator of sRNA biological relevance among the expanded candidate set recovered at RNIE score threshold 4. Across all nine species, 35.7% of sRNA candidates (750 of 2101) met the highly expressed criterion (mean depth ≥50 reads/position)

Mean coverage depth per candidate was computed over TSS- and RIT-delimited boundaries using samtools coverage, and a threshold of ≥50 reads/position was applied to define high-confidence transcriptional support. This threshold is absolute rather than dataset-normalized and should be interpreted accordingly:Datasets with the highest mean depth — *H. pylori* (2524 reads/position), *C. difficile* (1999 reads/position), *E. coli* (355 reads/position), and *S. enterica* (288 reads/position) —were dominated by candidates exceeding the ≥50 reads/position threshold. It should be noted, however, that this "highly expressed" category reflects sequencing coverage depth rather than intrinsic biological expression level, and varies across the RNA-seq projects analysed herecandidates. In contrast, low-depth datasets such as *S. oneidensis* (32 reads/position) and *M. extorquens* (41 reads/position) exhibited proportionally large weakly expressed fractions, suggesting that genuine sRNA transcripts were present but insufficiently covered to surpass classification thresholds a sequencing depth artifact rather than true absence of expression.

To evaluate condition-dependent sRNA expression, the pipeline was applied independently to five transcriptomic datasets from *Salmonella enterica* serovar Typhimurium ST4/74 (BioProject PRJNA215033; Kröger et al., 2013): mid-exponential phase (MEP), early stationary phase (ESP), late stationary phase (LSP), SPI-2-inducing conditions (InSPI2), and a pooled sample derived from 22 diverse environmental conditions (POOL).

Across all five conditions, 1225 unique loci were identified, revealing the breadth and plasticity of the *S. enterica* sRNA landscape. Of these, only 135 (11%) were shared across all five conditions simultaneously, defining a small constitutively expressed core. In stark contrast, 423 loci (35%) were strictly condition-specific — detected in exactly one condition and absent from all others demonstrating that the majority of the detectable sRNA repertoire is conditionally regulated rather than constitutively expressed. Among individual conditions, the number of detected loci varied substantially: POOL yielded the highest count (785), followed by ESP (669), InSPI2 (659), LSP (506), and MEP (443). The POOL dataset uniquely captured 104 loci undetected in any of the four individual conditions, reflecting aggregation of weak or transient transcriptional signals across 22 environments that fall below detection thresholds in any single profiled condition.

Condition-specific analysis revealed that ESP exhibited the highest number of unique candidates (165), substantially exceeding InSPI2 (47), LSP (66), MEP (41), and POOL (104). This enrichment of ESP-specific sRNAs is consistent with the extensive transcriptional reprogramming that accompanies the transition to nutrient limitation and aligns with the finding of Kröger et al. (2013) that ESP generates the highest number of unique transcriptions start sites among all profiled conditions reinforcing that transcription initiation is the primary determinant of condition-specific sRNA emergence.

The pooled dataset, despite maximizing overall detection sensitivity, paradoxically exhibited reduced condition-specificity relative to individual growth phases. The high locus count in POOL reflects cumulative signal aggregation across 22 environments rather than a coherent transcriptional program, complicating functional interpretation of candidates detected exclusively in this dataset. A single standard laboratory condition — the most common experimental design in non-model organisms captures between 36% (MEP: 443/1225) and 55% (ESP: 669/1225) of the total detectable repertoire, demonstrating that comprehensive sRNA discovery is structurally constrained by condition breadth. These findings underscore that multiplexed transcriptomics spanning diverse physiological states is not optional but essential for approaching exhaustive sRNA cataloguing in environmentally responsive organisms such as *S. enterica*.

### Limitations and Future Directions

Despite its robustness, several limitations constrain the pipeline’s scope. First, dRNA-seq data availability remains limited; organisms without these datasets cannot leverage TSS mapping unless an experimentally or computationally derived TSS list is available from other sources. Second, RIT detection assumes Rho-independent terminators predominate for sRNAs; organisms with Rho-dependent termination or atypical terminator structures may suffer systematic underestimation, as potentially illustrated by *C. jejuni* whose attenuated precision gain (9% final precision) despite a small initial prediction set suggests terminator detection limitations rather than expression-based noise. Third, condition sampling remains fundamental: even optimized pipelines cannot discover sRNAs unexpressed under profiled conditions, capping maximum completeness at approximately 40% for single growth condition strategies. Fourth, database gaps persist across BSRD, NCBI, Rfam, and RiboGap, conflating methodological precision with reference incompleteness and systematically underestimating true recall in understudied organisms.

Future sRNA discovery should pursue several complementary strategies. Multiplexed RNA-seq across diverse conditions — spanning nutritional, oxidative, osmotic, and host-mimicking stress contexts — would expand accessible repertoires and reveal condition-specific regulatory programs, as demonstrated by Kröger et al. 2013. Direct interaction mapping through iRIL-seq or CLIP-seq (Holmqvist et al. 2016) would provide orthogonal functional validation of predictions, independent of sequence homology. Computational tools designed to detect Rho-dependent termination signatures should be incorporated into future pipeline versions to address the systematic sRNA-knowledge gap in organisms where this termination mode is prevalent.

## Conclusion

This study presents a systematic framework for improving bacterial small regulatory RNA (sRNA) prediction through sequential integration of transcriptional context. By coupling sRNA detection with transcription start site (TSS) mapping and Rho-independent terminator (RIT) identification, we demonstrate substantial reductions in false-positive predictions while maintaining robust recovery of experimentally characterized sRNAs across phylogenetically diverse bacterial species.

The pipeline achieves 1.4- to 33-fold precision improvements across nine bacterial species, with initial candidate sets ranging from 319 to 29,947 reduced to biologically tractable lists of 22 to 809 prioritized candidates. A fundamental observation emerging from cross-species comparison is that baseline precision is primarily determined by the size of the initial prediction set rather than inherent biological differences between species. Organisms with compact initial prediction sets — H. pylori (536), C. jejuni (319), and C. difficile (452) — display higher baseline precisions (2.82–42.04%) simply because the ratio of known sRNAs to total predictions is more favorable. Conversely, species with large genomes or complex transcriptional backgrounds generating tens of thousands of predictions — M. extorquens (29,947), S. aureus (11,019), E. coli (9,335) — exhibit very low baseline precisions (0.03–1.96%) despite harboring equivalent or greater numbers of genuine sRNAs. This distinction is critical for correctly interpreting pipeline performance: fold-improvement is naturally larger for low-baseline species, while absolute final precision is naturally higher for compact-set species, but in both cases the pipeline consistently enriches true positives and eliminates noise.

Clostridioides difficile achieves the highest absolute final precision at 59% (22 final candidates from 452 initial predictions, 1.4-fold improvement), while M. extorquens DM4 achieves the greatest fold-improvement at 33-fold (112 final candidates from 29,947 initial predictions, 99.6% reduction), recovering 1 of 7 known sRNAs (14% recall). Staphylococcus aureus USA300 illustrates strong candidate reduction from 11,019 to 171 (98.4% reduction), recovering 5 of 88 known sRNAs (6% recall) at 3% final precision. Escherichia coli K-12 MG1655 recovered 38 of 116 known sRNAs (33% recall) at 7% final precision across 598 candidates, while Salmonella enterica ST4/74 recovered 48 of 143 (34% recall) at 7% final precision across 809 candidates — the highest recall values observed, consistent with their large and well-curated reference databases.

The robustness of TSS and RIT filtering is genome-size and phylogeny-independent, reflecting biological principles rather than organism-specific tuning. Sequential application of these filters preferentially eliminates transcriptional noise — degradation products, spurious initiation, and promoter read-through — while preserving true positives grounded in canonical promoter architecture and 3′ transcriptional termination.

Relaxation of RNIE bit score thresholds from default parameters to -T 4 recovered previously undetected sRNAs, including the experimentally validated sRNA1969 in M. extorquens, validating the hypothesis that standardized terminator models may miss species-specific variants. Such calibration may be a necessary compromise for non-model organisms underrepresented in terminator model training datasets, extending pipeline applicability across bacterial phylogeny.

Precision variation across species correlates with both initial prediction set size and reference annotation completeness rather than methodological failure. H. pylori (81 BSRD sRNAs), E. coli (116 sRNAs), S. enterica (143 sRNAs), and S. aureus (88 SRD sRNAs) have received decades of transcriptomic investigation providing dense reference sets, while M. extorquens (7 RiboGap entries), C. jejuni (22 sRNAs), S. oneidensis (17 sRNAs), and B. amyloliquefaciens (28 BSRD entries) represent understudied genomes where reference incompleteness systematically underestimates true recall. Lower apparent precision in these organisms likely reflects pipeline recovery of genuine sRNAs absent from incomplete reference sets. This distinction is critical: for organisms with sparse annotation, unmatched predictions represent opportunities for new sRNA gene discovery rather than evidence of pipeline failure.

RNA-seq expression depth emerged as a reliable secondary indicator of sRNA biological relevance, though its interpretation is dataset-dependent. Candidates supported by ≥50 reads per base exhibit consistent expression above noise thresholds, while cross-species comparisons require accounting for sequencing depth variation across projects. Multi-condition analysis of S. enterica across five conditions revealed that only 135 of 1225 unique loci (11%) were detected across all conditions simultaneously, while 423 loci (35%) were strictly condition-specific, underscoring that comprehensive sRNA discovery requires multiplexed transcriptomics spanning diverse physiological contexts — a structural constraint inherent to all prediction pipelines regardless of algorithmic sophistication.

The integration of TSS and RIT constraints directly addresses documented limitations of expression-based tools: sRNA-Detect alone predicts 7,812 transcripts in S. enterica at 1.42% baseline precision (111 known sRNA matches). Our pipeline maintained 34% recall while reducing final candidates to 809, rendering high-throughput experimental validation tractable. Comparative benchmarking against published tools confirms that systematic incorporation of transcriptional context markers outperforms sensitivity-focused alternatives, achieving superior precision-recall trade-offs particularly for non-model organisms.

However, the pipeline’s scope is constrained by fundamental limitations. dRNA-seq data availability remains restricted, and organisms relying predominantly on Rho-dependent termination — potentially including C. jejuni — exhibit reduced terminator recovery density. sRNAs derived from post-transcriptional processing, including tRNA-derived fragments and UTR-derived RNAs, lack independent TSS and RIT signatures and remain undetectable by this framework. Condition-specific expression caps theoretical maximum single-condition discovery rates at approximately 40%.

Future efforts should pursue: (1) Multiplexed RNA-seq across diverse conditions to expand accessible sRNA repertoires; (2) Direct interaction mapping via iRIL-seq or CLIP-seq for orthogonal functional validation; (3) Incorporation of Rho-dependent termination detection; (4) Integration of machine learning classifiers trained on experimentally validated sRNAs; and (5) Target prediction coupled to candidate assessment to reveal biological relevance independently of sequence homology.

In conclusion, transcriptional context markers (TSS and RIT) constitute universal, sequence-independent biological filters applicable across bacterial phylogeny. Their sequential integration substantially enriches sRNA prediction precision while maintaining sensitivity for genuine regulatory molecules, providing a generalizable framework for improving bacterial RNA annotation and discovering novel regulatory elements in understudied organisms where dense reference annotation remains inaccessible.

## ACKNOWLEDGMENTS

This research was enabled by computational resources provided by the Digital Research Alliance of Canada (formerly Compute Canada). This research was in part funded by an NSERC discovery grant (RGPIN-2025-06433) Fondation Armand-Frappier and CRIPA.

## AUTHOR CONTRIBUTIONS

Sinda Elhedi: pipeline design, data analysis, writing. Kadidia Dite Selly N’Diaye: Contributing with data of *Methylorubrum extorquens*,J. Perreault: Supervision, writing – review and editing.

## DATA AVAILABILITY

All RNA-seq datasets used in this study are publicly available through NCBI GEO and ENA repositories; accession numbers are provided in Table. Script and databases are available on GitHub : https://github.com/sindaelhedi/sRNA_TSS_RIT_pipeline.git

