## Supplementary for "A pipeline for identifying small noncoding RNA (sRNA) candidates in bacteria"

**[a] INRS – Armand-Frappier Health Biotechnology Centre, 531 boulevard des Prairies, Laval, QC, H7V 1B7, Canada,**

### Figures

Figure S1:Stringency Analysis Grid ........................................................ 3
Figure S2:Effect of score threshold on sRNA prediction across bacterial species........................................................................................ 6

### Tables

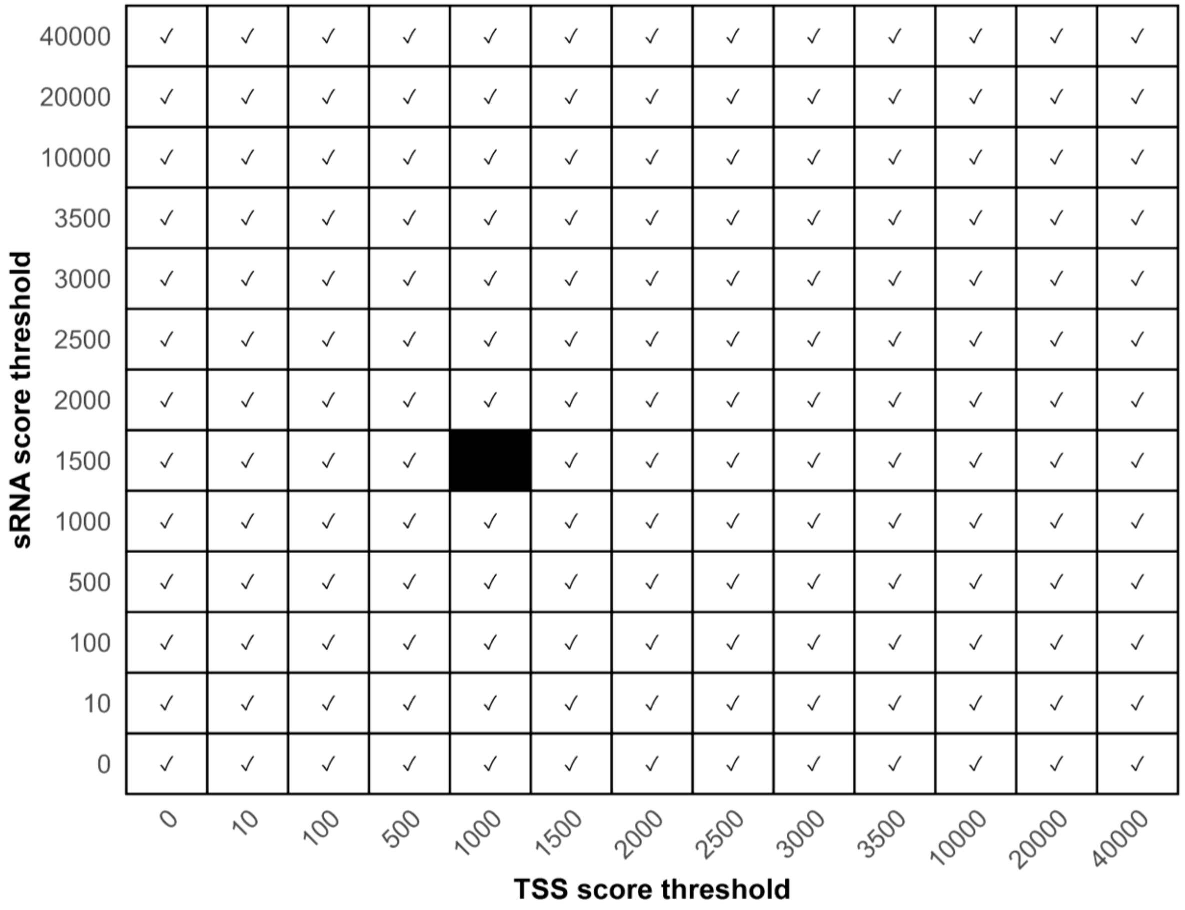

Figure S1:Stringency Analysis Grid

*Each cell represents a unique combination of TSS score and sRNA score thresholds. A total of 169 threshold combinations were evaluated for each species. One example combination (TSS = 1000, sRNA = 1500) is highlighted to illustrate how parameter pairs define a specific stringency condition.*

| Category | Species | Strain | Genome size (Mb) | BioProject accession | Experimental conditions | Ref. |
|---|---|---|---|---|---|---|
| Benchmark | *Escherichia coli* | K-12 MG1655 | 4.64 | PRJNA238884 | LB, minimal medium; exponential/stationary phases | Thomason et al.2015 |
| | *Staphylococcus aureus* | USA300 | 2.90 | PRJEB23980 | Various antibiotic treatments | Choe et al. 2018 |
| | *Salmonella enterica* | Typhimurium ST4/74 | 4.88 | PRJNA215033 | 22 distinct growth conditions | Kröger et al. 2013 |
| Test | *Bacillus amyloliquefaciens* | FZB42 | 3.92 | PRJNA277720 | Sporulation, biofilm formation | Fan et al. 2015 |
| | *Helicobacter pylori* | 26695 | 1.67 | PRJNA280272 | Rich medium, stationary phase | Bischler et al. 2015 |
| | *Campylobacter jejuni* | 81116 | 1.63 | PRJNA169137 | Rich medium, multiple growth conditions | Dugar G et al 2013 |
| | *Methylorubrum extorquens* | DM4 | 6.12 | PRJEB52788 | Dichloromethane, methanol (carbon sources) | Maucourt, B., et al. 2022 |
| | *Shewanella oneidensis* | MR-1 | 4.97 | PRJNA251904 | LB, defined lactate minimal medium | Shao et al. 2014 |
| | *Clostridium difficile* | 630Δerm | 4.29 | PRJNA626554 | Exponential phase (4h), stationary phase (10h), nutrient starvation | Soutourina et al. 2020 |

Table S1:Benchmark and test bacterial species used in this study.

*The table summarizes the species, strains, genome sizes, BioProject accessions, and experimental conditions associated with RNA-seq datasets used for sRNA detection and validation. Benchmark species correspond to well-characterized organisms with extensive annotated sRNA repertoires, while test species represent less-characterized organisms.*

| ***Species*** | ***Known sRNAs*** | ***Initial candidates (sRNA-Detect)*** | ***predicted sRNA that hit known sRNA(bit score>60)*** | ***Baseline precision*** | ***+ TSS (candidates)*** | ***predicted sRNA that hit known sRNA*** | ***+ TSS precision*** | ***TSS+RIT candidates*** | ***predicted sRNA that hit known sRNA*** | ***+TSS+ RIT precision*** | ***Recovered final known sRNAs*** | ***Recall*** |
|---|---|---|---|---|---|---|---|---|---|---|---|---|
| ***S. aureus*** | ***88*** | ***11 019*** | ***131*** | ***1,19%*** | ***1 594*** | ***21*** | ***1,30%*** | ***171*** | ***5*** | ***3%*** | ***5*** | ***6%*** |
| ***E. coli*** | ***116*** | ***9 335*** | ***183*** | ***1,96%*** | ***3 598*** | ***104*** | ***2,90%*** | ***598*** | ***41*** | ***7%*** | ***38*** | ***33%*** |
| ***S. enterica*** | ***143*** | ***7 812*** | ***111*** | ***1,42%*** | ***5 465*** | ***75*** | ***1,40%*** | ***809*** | ***57*** | ***7%*** | ***48*** | ***34%*** |
| ***B. amyloliquefaciens*** | ***28*** | ***806*** | ***4*** | ***0,50%*** | ***492*** | ***3*** | ***0,60%*** | ***160*** | ***3*** | ***2%*** | ***3*** | ***11%*** |
| ***M. extorquens*** | ***7*** | ***29 947*** | ***8*** | ***0,03%*** | ***4 029*** | ***1*** | ***0,00%*** | ***112*** | ***1*** | ***1%*** | ***1*** | ***14%*** |
| ***S. oneidensis*** | ***17*** | ***6 137*** | ***12*** | ***0,20%*** | ***1 819*** | ***5*** | ***0,30%*** | ***163*** | ***2*** | ***1%*** | ***2*** | ***12%*** |
| ***H. pylori*** | ***81*** | ***536*** | ***31*** | ***5,78%*** | ***128*** | ***20*** | ***15,60%*** | ***32*** | ***7*** | ***22%*** | ***6*** | ***7%*** |
| ***C. jejuni*** | ***22*** | ***319*** | ***9*** | ***2,82%*** | ***158*** | ***8*** | ***5,10%*** | ***34*** | ***3*** | ***9%*** | ***3*** | ***14%*** |
| ***C. difficile*** | ***14*** | ***452*** | ***190*** | ***42,04%*** | ***132*** | ***60*** | ***45,50%*** | ***22*** | ***13*** | ***59%*** | ***2*** | ***14%*** |

Table S2:Progressive precision improvements accross pipeline stages.

Pipeline performance across nine phylogenetically diverse bacterial species. Initial candidates: initial sRNA-Detect predictions;+ TSS: candidates intersecting with transcription start sites (±60/-20 bp); +**TSS+RIT**: candidates with TSS and downstream Rho-independent terminators. n: number of candidates; Hits: predicted sRNAs match to known sRNAs databases; %: precision (Hits/n × 100); Recall: fraction of known sRNAs recovered (comparing to known sRNAs).

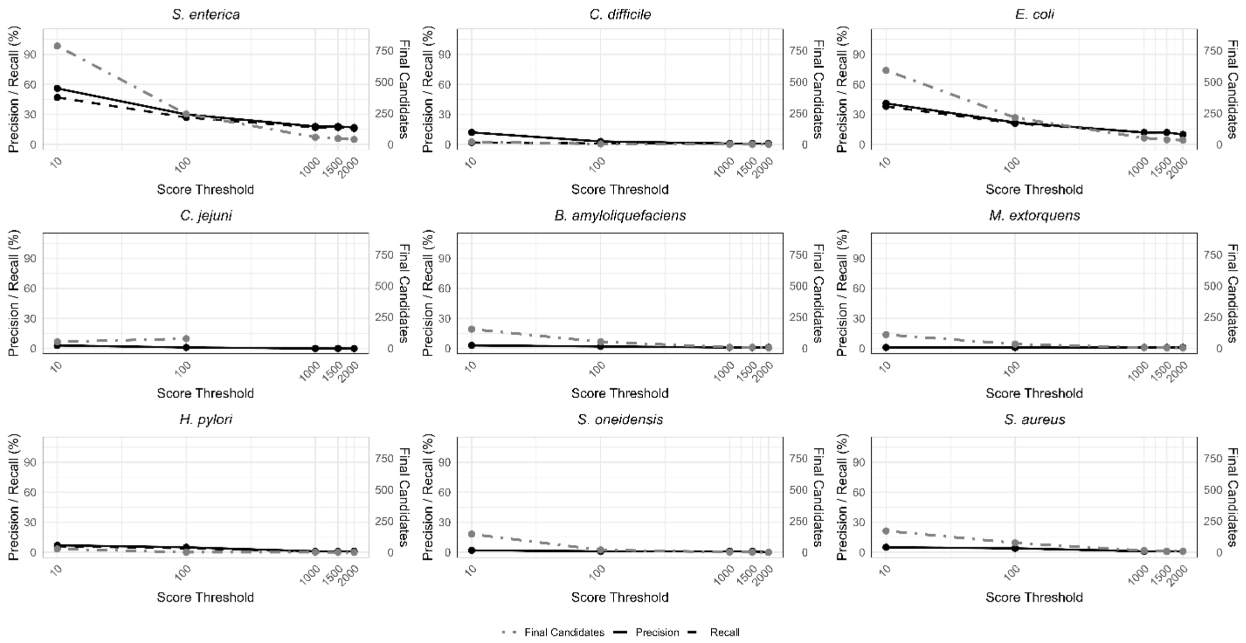

Figure S2:Effect of score threshold on sRNA prediction across bacterial species

Effect of score threshold on sRNA prediction performance across nine bacterial species. Precision (solid line) and recall (dashed line) are shown on the primary y-axis, while the number of final candidates (dot-dashed line) is shown on the secondary y-axis.

| | | | | |
|---|---|---|---|---|
| NC_009839.1 | 100737 | 100833 | CJnc10 | + |
| NC_009839.1 | 248101 | 248257 | CJnc20 | - |
| NC_009839.1 | 518347 | 518664 | RnpB | - |
| NC_009839.1 | 681024 | 681305 | Cjnc60 | - |
| NC_009839.1 | 879954 | 880052 | CJnc80 | - |
| NC_009839.1 | 996242 | 996319 | CJnc100 | + |
| NC_009839.1 | 1148712 | 1148849 | CJnc110 | + |
| NC_009839.1 | 1200515 | 1200700 | 6S | + |
| NC_009839.1 | 1209853 | 1209926 | CJnc140 | + |
| NC_009839.1 | 1441288 | 1441362 | TracrRNA | + |
| NC_009839.1 | 1563091 | 1563246 | CJnc180 | + |
| NC_009839.1 | 1563120 | 1563337 | CJnc190 | - |
| NC_009839.1 | 1624612 | 1624711 | CJnc230 | - |
| NC_009839.1 | 543100 | 543269 | CJnc21 | + |
| NC_009839.1 | 591312 | 591426 | CJnc30 | + |
| NC_009839.1 | 1455231 | 1455269 | crRNA2 | + |
| NC_009839.1 | 1455362 | 1455400 | crRNA4 | + |
| NC_009839.1 | 1559675 | 1559722 | CJnc170 | + |
| NC_009839.1 | 1589597 | 1589666 | CJas_Cj1667c | + |
| NC_009839.1 | 1152713 | 1152810 | CJnc120 | + |
| NC_009839.1 | 75877 | 75984 | SRP | + |
| NC_009839.1* | 1626903 | 1626741 | HPnc0260 | - |

Table S3:List of known sRNAs of campylobacter jejuni recovered from literature.

Most of the annotated sRNAs originate from the study by Dugar *et al.* (2013), and only those experimentally validated by Northern blot were retained. *HPnc0260* was obtained from the Rfam database.

| *Species* | *Known sRNAs* | *Initial candidates (sRNA-Detect)* | *Baseline precision* | *After TSS (candidates)* | *After TSS (precision %)* | *TSS+RIT candidates (score -T 4)* | *After RIT (precision %)* | *Matched sRNAs count* | *Recall* |
|---|---|---|---|---|---|---|---|---|---|
| *S. aureus* | *88* | *11019* | *97.4%* | *1 594* | *98%* | *171* | *98.8%* | *87* | *98.8%* |
| *E. Coli* | *116* | *9335* | *25%* | *3 598* | *99.6%* | *598* | *99.5%* | *79* | *65.3%* |
| *S. Enterica* | *143* | *10595* | *18.3%* | *5 586* | *22.2%* | *779* | *25.9%* | *101* | *70.6%* |
| *Bacillus* | *28* | *806* | *11.9%* | *492* | *14.8%* | *160* | *22.5%* | *15* | *53.5%* |
| *M. Extorquens* | *7* | *29947* | *25%* | *4 029* | *30.3%* | *112* | *30.4%* | *6* | *85.7%* |
| *S. Oneidensis* | *17* | *6 137* | *1.3%* | *1819* | *14.5%* | *163* | *7.97%* | *4* | *24%* |
| *H. Pylori* | *81* | *536* | *45.3%* | *128* | *53.9%* | *32* | *62.5%* | *23* | *28.4%* |
| *C. Jejuni* | *22* | *319* | *49%* | *158* | *43.7%* | *34* | *52.9%* | *14* | *63%* |
| *C. Difficile* | *14* | *452* | *54.6%* | *132* | *92.4%* | *22* | *86%* | *4* | *29%* |

Table S4:Progressive precision improvements across pipeline stages with permissive blast parameters.

Pipeline performance across nine phylogenetically diverse bacterial species. Initial candidates: initial sRNA-Detect predictions; + TSS: candidates intersecting with transcription start sites (±60/-20 bp); +TSS+RIT: candidates with TSS and downstream Rho-independent terminators. n: number of candidates; Hits: predicted sRNAs match to known sRNAs databases; %: precision (Hits/n × 100); Recall: fraction of known sRNAs recovered (comparing to known sRNAs). These results were obtained for permissive blast parameters without bit score filters.
